# Mass Recovery following Caloric Restriction Reverses Lipolysis and Proteolysis, but not Gluconeogenesis, in Insulin Resistant OLETF Rats

**DOI:** 10.1101/2021.05.17.444436

**Authors:** Manuel A. Cornejo, Jaapna Dhillon, Akira Nishiyama, Daisuke Nakano, Rudy M. Ortiz

## Abstract

Caloric restriction (CR) is one of the most important behavioral interventions to reduce excessive abdominal adiposity, which is a risk factor for the development of insulin resistance. Previous metabolomics studies have characterized substrate metabolism during healthy conditions; however, the effects of CR and subsequent mass recovery on shifts in substrate metabolism during insulin resistance (IR) have not been widely investigated. To assess the effects of acute CR and the subsequent mass recovery on shifts in substrate metabolism, a cohort of 15-week old Long Evans Tokushima Otsuka (LETO) and Otsuka Long Evans Tokushima Fatty (OLETF) rats were calorie restricted (CR: 50% × 10 days) with or without partial body mass recovery (PR; 73% x 7 days), along with their respective *ad libitum* controls. End-of-study plasma samples were analyzed for primary carbon metabolites by gas chromatography (GC) time-of-flight (TOF) mass spectrometry (MS) data acquisition. Data analysis included PCA, Pearson correlation vs previously reported variables (adipose and body masses, and insulin resistance index, IRI), and metabolomics maps (MetaMapp) generated for the most significant group comparisons. All treatments elicited a significant group differentiation in at least one principal component. CR improved TCA cycle in OLETF, and increased lipolysis and proteolysis. These changes were reversed after PR except for gluconeogenesis. Plasma lipid concentrations were inversely correlated to IRI in LETO, but not OLETF. These shifts in substrate metabolism suggest that the CR-induced decreases in adipose may not be sufficient to more permanently alter substrate metabolism to improve IR status during metabolic syndrome.

## Introduction

Abdominal adiposity, concomitant with increased plasma leptin and branched chain amino acids (BCAA), and lower adiponectin, are risk factors for insulin resistance (IR) independent of higher protein intake and body mass index (BMI) [1, 2]. Also, obesity leads to excessive β-oxidation (e.g. mitochondrial overload) in skeletal muscle concomitant with decreased fatty acid oxidation in liver, which in turn may be responsible for muscle insulin resistance [3]. Excessive β-oxidation is also associated with an increase of glucose-6-phosphate dehydrogenase (G6PD), the rate limiting enzyme in the pentose phosphate pathway (PPP), which in turn increases circulating triglyceride (TG), non-esterified fatty acids (NEFA), and pro inflammatory cytokines [4]. Moreover, increased BCAA, and glucuronic and hexuronic acids are correlated with the progression of type 2 diabetes mellitus (T2DM) in rats [5].

Caloric restriction (CR), especially when associated with a decrease in carbohydrate intake [6, 7], is an important behavioral intervention to reduce excess adipose mass [8]. In healthy mice, short term CR (30% for 16 days) increased the hepatic glycogenic amino acid, valine (a BCAA), and lean tissue turnover, and decreased fasting glucose and VLDL [9]. In healthy humans, acute, severe CR (90%, 48 hrs) decreased glucose and pyruvate, and increased glucogenic amino acids, ketone bodies, and lipolysis products, but these changes were reversed after a 48-h *ad libitum* refeeding [10] demonstrating the shifts in critical metabolic pathways to acute alterations in caloric intake. In elderly patients, enteral refeeding after 10 days of food deprivation resulted in a decrease in acylcarnitines concomitant with an increase of free amino acids and higher urea cycle activity [11]. However, to the best of our knowledge there is still little information on the shifts in biochemical processes (assessed via metabolomics) in response to CR and the subsequent mass recovery from refeeding during metabolic syndrome.

The OLETF rat develops early onset hyperglycemia, obesity, insulin resistance, hyperlipidemia, and hypertension, all components of the metabolic syndrome [12-15]. Moreover, OLETF rats develop an age-associated reduction in hepatic beta-hydroxyacyl-coA dehydrogenase (β-HAD) and citrate synthase, which are rate limiting steps of β-oxidation and tricarboxylic acid cycle (TCA) cycle, respectively [16]. In addition, OLETF rats present with lower plasma levels of tryptophan and its metabolite, kynurenine, compared with their healthy, lean strain control Long-Evans Tokushima Otsuka (LETO). These variables may be predictive markers for the development of T2DM [17]. We hypothesized that reducing abdominal adiposity (i.e. visceral adipose) during metabolic syndrome via an acute CR without a modification in the macronutrient proportion, reduces gluconeogenesis, the pentose phosphate pathway (PPP), and plasma BCAA concentrations, with the advantage of increasing the rate of TCA cycle, lipolysis, and β-oxidation, without compromising lean tissue catabolism (lean tissue proteolysis). However, based on our previous results, we also hypothesized that partial recovery of body mass, even without full adipose recovery, is sufficient to fully revert these changes.

## Method

Details of the current study have been published previously [18] and summarized in **Figure 1**. The current study complements the previous data by using metabolomics approaches to examine the shifts in metabolism associated with changes in body mass induced by CR and refeeding. The study was approved by IACUC of Kagawa Medical University.

**Figure 1.**
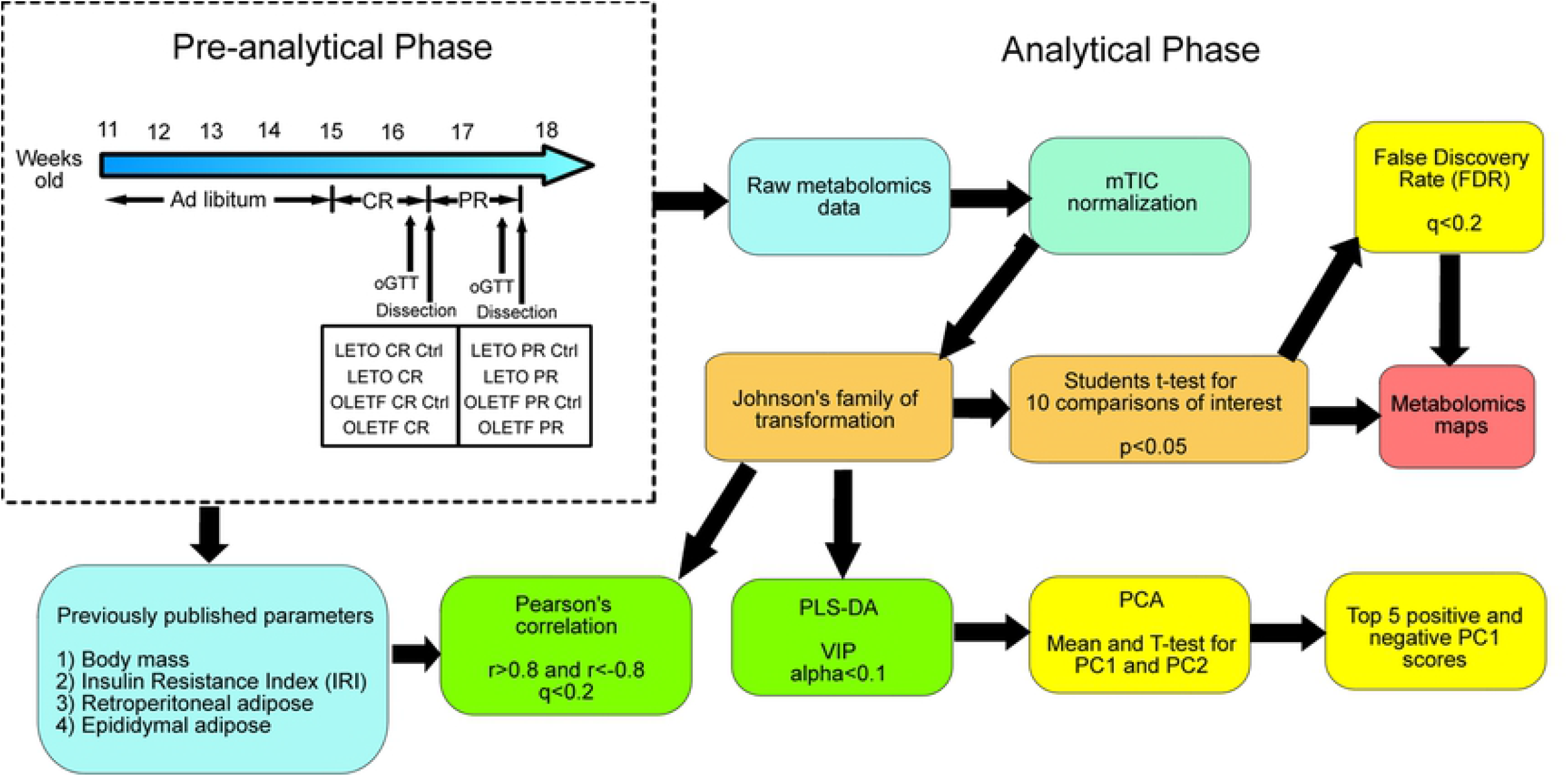
Project flowchart. CR: Caloric Restriction; LETO: Long-Evans Tokushima Otsuka; oGTT: oral Glucose Tolerance Test; OLETF: Otsuka Long-Evans Tokushima Fatty; PCA: Principal Component Analysis; PLS-DA: Partial Least Squares Discriminant Analysis; PR: Partial Recovery; VIP: Variable Importance in the Projection.

### Caloric Restriction (CR) and Partial Recovery (PR)

Briefly, lean strain control LETO (n=29) and obese, insulin resistant OLETF rats (n=29) were fed *ad libitum* with standard laboratory rat chow (MF; Oriental Yeast Corp., Tokyo, Japan) for 4 weeks. At 15 weeks of age, rats were separated in two *ad libitum* food control groups (n=8/group/strain), two CR groups (n=7/group/strain), and two partial recovery (PR) groups (n=7/group/strain). Both CR and PR groups were subjected to 50% CR (compared to *ad libitum* control) for 10 days. CR groups were subjected to an oral glucose tolerance test (oGTT) and dissected three days later along with their respective *ad libitum* controls (CR Ctrl, n=8/group/strain). Meanwhile, PR groups were fed *ad libitum* for 7 days, achieving partial body mass recovery (73% recovery of mass loss) before an oGTT and dissection paired with their PR control groups (n=7/group/strain). Both CR and PR control groups were fed *ad libitum* for the entire study. However, they were considered independent groups to avoid the confounding factor of age as they were dissected 1 week apart; however, no significant differences were detected between CR and PR control groups despite the slight difference in age suggesting that age was not a confounding factor here. Animals were maintained in a specific pathogen-free facility under controlled temperature (23°C) and humidity (55%) with a 12-h light, 12-h dark cycle. All animals were given free access to water for the entire study.

### Oral Glucose Tolerance Test (oGTT)

Oral glucose tolerance tests (oGTT) were performed as previously reported [18] and insulin resistance index (IRI) was calculated as previously described [13] and reported [18]. Data were used here to correlate the shifts in metabolites with IRI.

### Blood Sample and Tissue Collection

Details of blood sample and tissue collections have been previously reported [18]. Briefly, animals were anesthetized after an overnight fasting with a 100 mg/kg i.p. pentobarbital injection and retroperitoneal and epididymal fat depots were dissected, weighed, and collected for other analyses. Arterial blood was collected via the abdominal aorta into chilled vials containing a cocktail of 50 mmol/L EDTA, 5000 KIU aprotinin, and 0.1 mmol sitagliptin phosphate (DPP4 inhibitor). Blood samples were centrifuged (3,000 g, 15min at 4°C), and the plasma was transferred to cryo-vials and immediately stored at −80°C. Aliquots of plasma (n=5 per group/strain) were analyzed for primary carbon metabolites by gas chromatography (GC) time-of-flight (TOF) mass spectrometry (MS) data acquisition and processing at the West Coast Metabolomics Center as previously described [19], generating a dataset of 143 consistently identified metabolites.

### Data Analysis

#### Data Normalization and Transformation

Data were reported as quantitative ion peak heights and were normalized by the sum peak height of all structurally annotated compounds (mTIC normalization) [20]. Transformations were done using Johnson’s family of transformation, depending on data normality, as described in other studies [21]. Student’s t-test was performed in 10 previously selected pairwise comparisons (**Table 1**) and Benjamini-Hochberg False Discovery Rate (FDR) correction [22] q values were calculated for each metabolite per comparison.

**Table 1.**
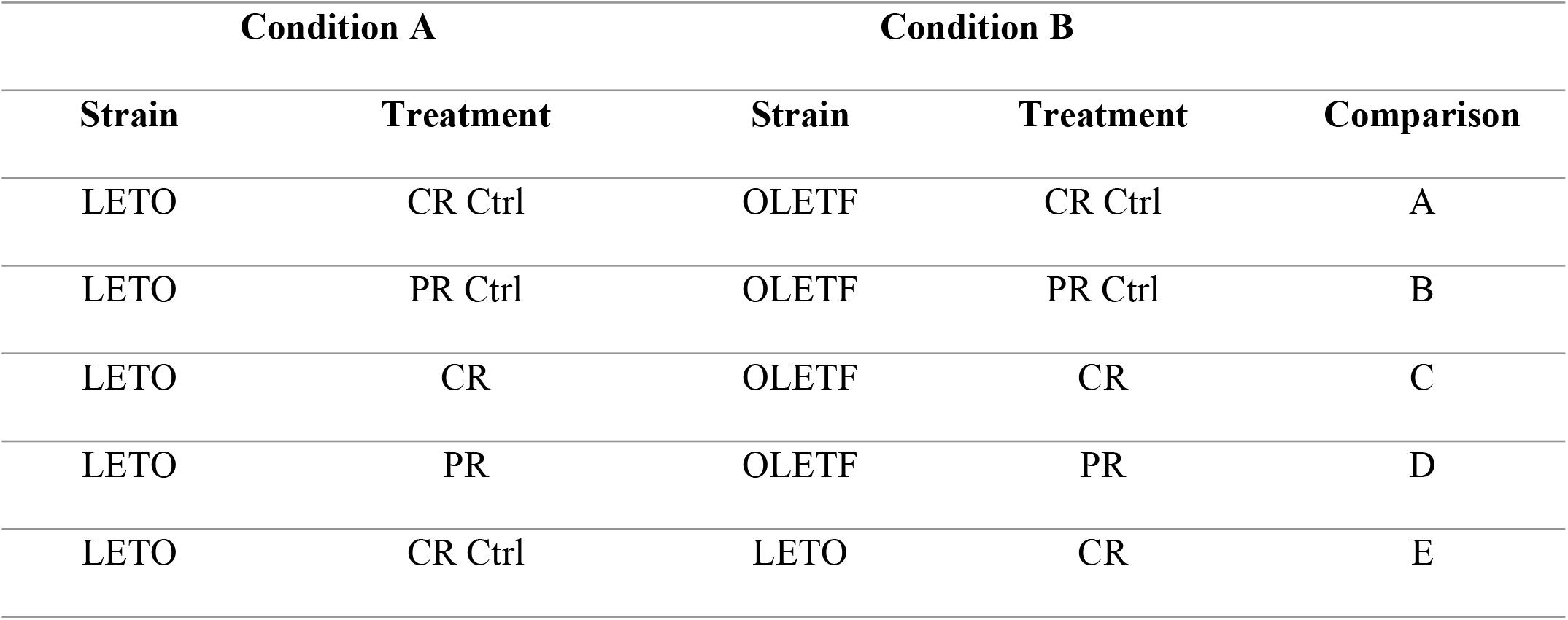
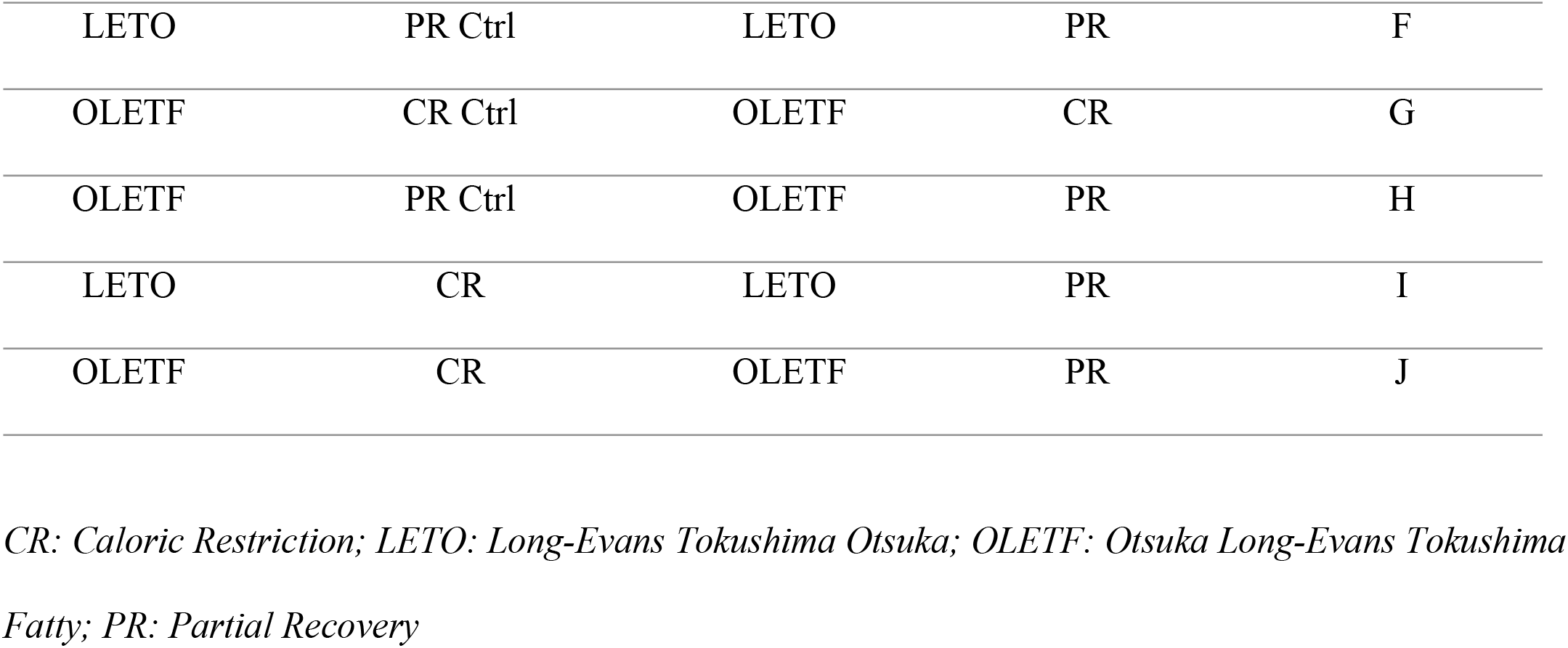
Fold changes were calculated as mean condition B / mean condition A, after mTIC normalization.

| Condition A |  | Condition B |  | Comparison |
| --- | --- | --- | --- | --- |
| Strain | Treatment | Strain | Treatment |  |
| LETO | CR Ctrl | OLETF | CR Ctrl | A |
| LETO | PR Ctrl | OLETF | PR Ctrl | B |
| LETO | CR | OLETF | CR | C |
| LETO | PR | OLETF | PR | D |
| LETO | CR Ctrl | LETO | CR | E |
| LETO | PR Ctrl | LETO | PR | F |
| OLETEF | CR Ctrl | OLETEF | CR | G |
| OLETEF | PR Ctrl | OLETEF | PR | H |
| LETO | CR | LETO | PR | I |
| OLETEF | CR | OLETEF | PR | J |

CR: Caloric Restriction; LETO: Long-Evans Tokushima Otsuka; OLETEF: Otsuka Long-Evans Tokushima Fatty; PR: Partial Recovery

#### Metabolic Mapping

Fold changes were calculated as mean condition B / mean condition A, after mTIC normalization. The conditions are described in **Table 1**. Metabolic maps were plotted per comparison only for metabolites with p<0.05 after Student’s t-test using MetaMapp [23] and visualized in Cytoscape 3.7.1 with organic layout. Nodes with significant difference after FDR correction (q<0.2) were highlighted for fold-change and direction of change. Metabolites were classified into either primary or secondary metabolism pathways based on their entry on the Kyoto Encyclopedia of Genes and Genomes (KEGG) database [24], where secondary metabolism encompasses either cofactor, vitamin or xenobiotic metabolism. The maps with the most significant differences are shown in **Figure 2** and further discussed.

**Figure 2.**
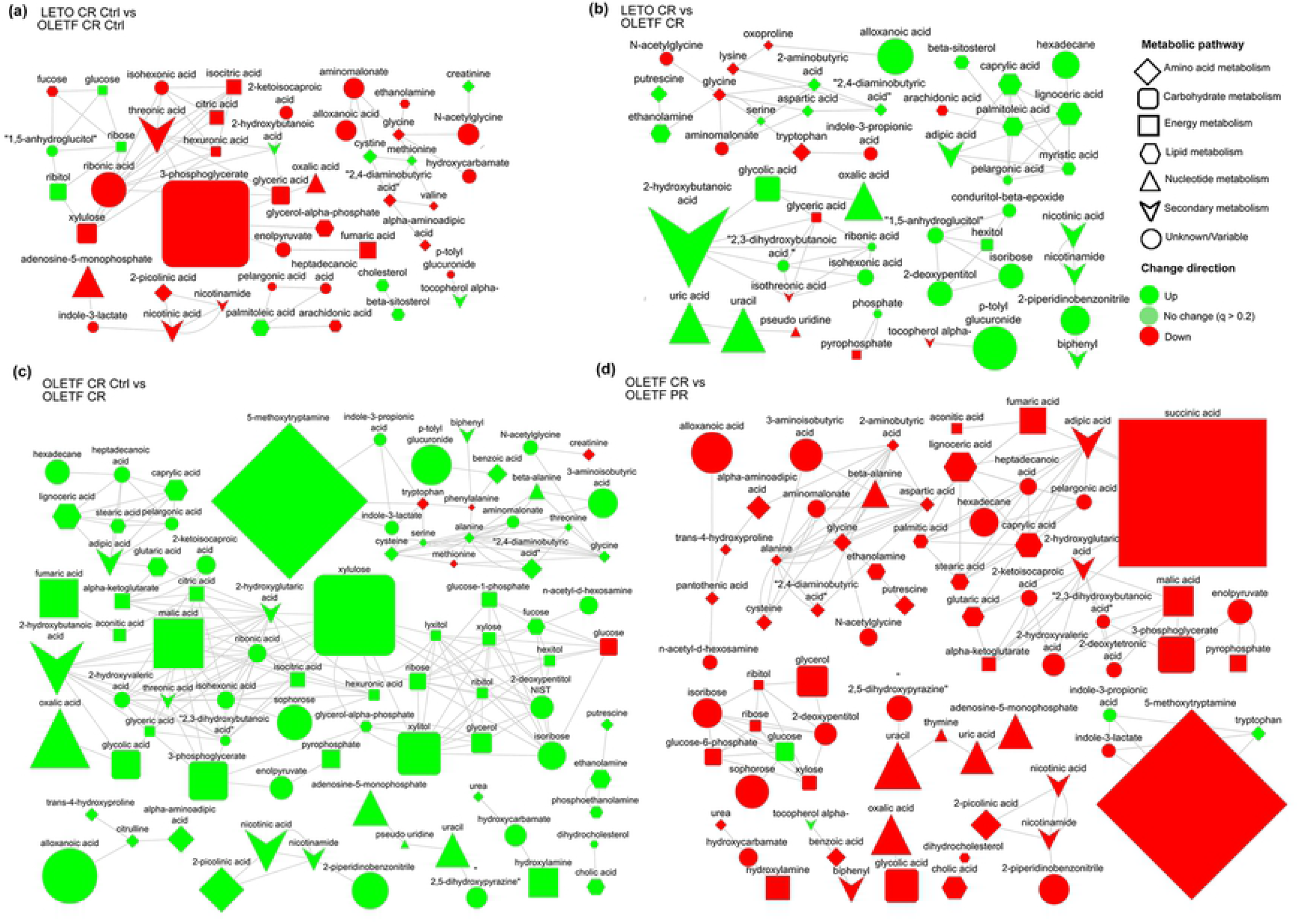
Metabolomics maps showing metabolites with p<0.05 after Student’s t-test, representing mean plasma peak intensity fold-changes between groups in metabolites with q<0.2, for (a) LETO CR Ctrl vs OLETF CR Ctrl (Comparison A), (b) LETO CR vs OLETF CR (Comparison C), (c) OLETF CR Ctrl vs OLETF CR (Comparison G), (d) OLETF PR Ctrl vs OLETF PR (Comparison J).

#### Correlations with Body Mass, Insulin Resistance Index, and Adipose Depots

Pearson correlations were calculated for parameters previously reported [18]. Johnson transformed body mass, IRI, and retroperitoneal and epididymal adipose masses were treated as independent variables and Johnson transformed metabolites were treated as dependent variables. Correlations were considered strong at r<-0.8 or r>0.8 and q<0.2. Statistical analyses were performed in JMP pro (version 14, SAS Institute Inc., Cary, NC, USA).

#### Partial Least Squares-Discriminant Analysis (PLS-DA) and Principal Component Analysis (PCA)

PLS-DA was run on all metabolites for all 10 comparisons (**Table 1**) by nonlinear iterative partial least squares (NIPALS), with leave-one-out validation, four (n-1) factors and variable importance in the projection (VIP) score threshold (alpha) of 0.1. The metabolites above the threshold were used further for PCA for correlation analyses. The scripts showing the metabolites considered for each PCA analysis can be found in the online repository. The rationale behind using PCA after PLS-DA was to avoid the increased group separation due to artifacts inherent to PLS-DA, and instead visually reflect the variability that distinguishes the groups [25]. For each PCA, mean of scores for PC1 and PC2, and standard deviations were calculated, and a t-test performed for both PC for significant difference (*P*<0.05) (**Table 2**). For each comparison, the metabolites with the five highest positive and 5 highest negative loadings are shown in **Table 3**.

**Table 2.** Mean PC score ± SD for PC1 and PC2 for each group comparison and T-test P value for each comparison.

| Comparison | Group | Mean PC1 | P value | Mean PC2 | P value |
| --- | --- | --- | --- | --- | --- |
| **A** | LETO CR Ctrl | 5.16 ± 3.58 | 0.0019 | -1.00 ± 3.76 | 0.3091 |
| | OETF CR Ctrl | -5.16 ± 1.15 | | 1.00 ± 1.17 | |
| **B** | LETO PR Ctrl | 5.06 ± 1.50 | <.0001 | -0.38 ± 2.59 | 0.6400 |
| | OETF PR Ctrl | -5.06 ± 1.13 | | 0.38 ± 2.41 | |
| **C** | LETO CR | -3.32 ± 0.76 | 0.0622 | -1.72 ± 0.23 | 0.1093 |
| | OETF CR | 3.32 ± 5.80 | | 1.72 ± 3.74 | |
| **D** | LETO PR | 5.01 ± 1.82 | <.0001 | -0.12 ± 3.39 | 0.8859 |
| | OETF PR | -5.01 ± 1.04 | | 0.12 ± 1.52 | |
| **E** | LETO CR Ctrl | -4.14 ± 0.47 | 0.0105 | 0.88 ± 3.42 | 0.3640 |
| | LETO CR | 4.14 ± 4.13 | | -0.88 ± 2.20 | |
| **F** | LETO PR Ctrl | 4.48 ± 1.85 | <.0001 | -0.31 ± 3.60 | 0.7775 |
| | LETO PR | -4.48 ± 1.11 | | 0.31 ± 3.09 | |
| **G** | OETF CR Ctrl | -4.65 ± 0.51 | 0.0158 | -0.36 ± 0.32 | 0.7729 |
| | OETF CR | 4.65 ± 5.20 | | 0.36 ± 5.17 | |
| **H** | OETF PR Ctrl | 3.58 ± 3.89 | 0.0139 | 1.42 ± 3.46 | 0.1457 |
| | OETF PR | -3.58 ± 0.57 | | -1.42 ± 1.33 | |
| **I** | LETO CR | 4.51 ± 2.36 | 0.0003 | -0.40 ± 2.84 | 0.6334 |
| | LETO PR | -4.51 ± 1.07 | | 0.40 ± 2.16 | |
| **J** | OETF CR | 4.28 ± 5.18 | 0.0207 | 0.22 ± 5.20 | 0.8595 |
| | OETF PR | -4.28 ± 0.44 | | -0.22 ± 0.46 | |

CR: Caloric Restriction; LETO: Long-Evans Tokushima Otsuka; OETF: Otsuka Long-Evans Tokushima Fatty; PC: Principal Component; PR: Partial Recovery

**Table 3.**
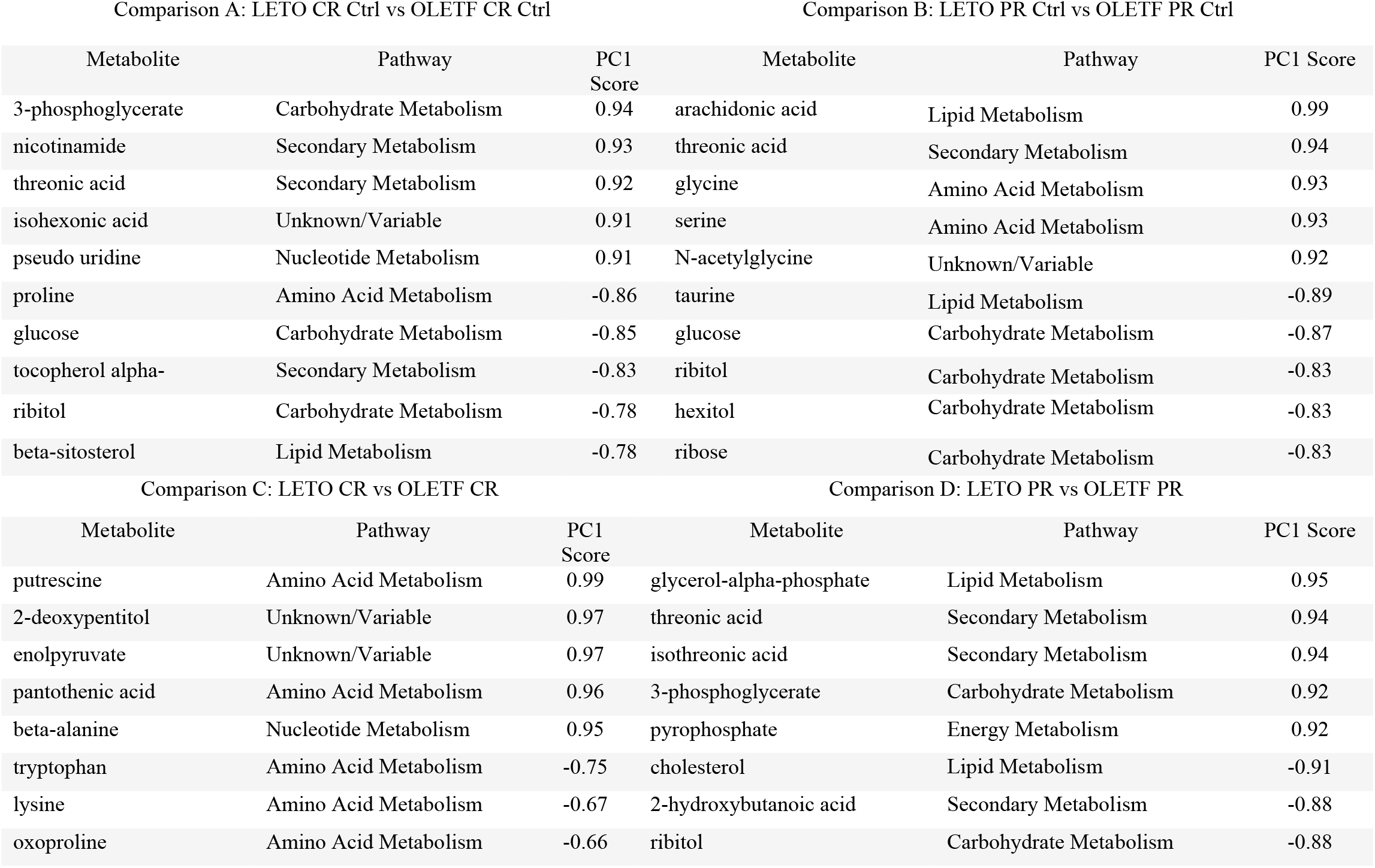
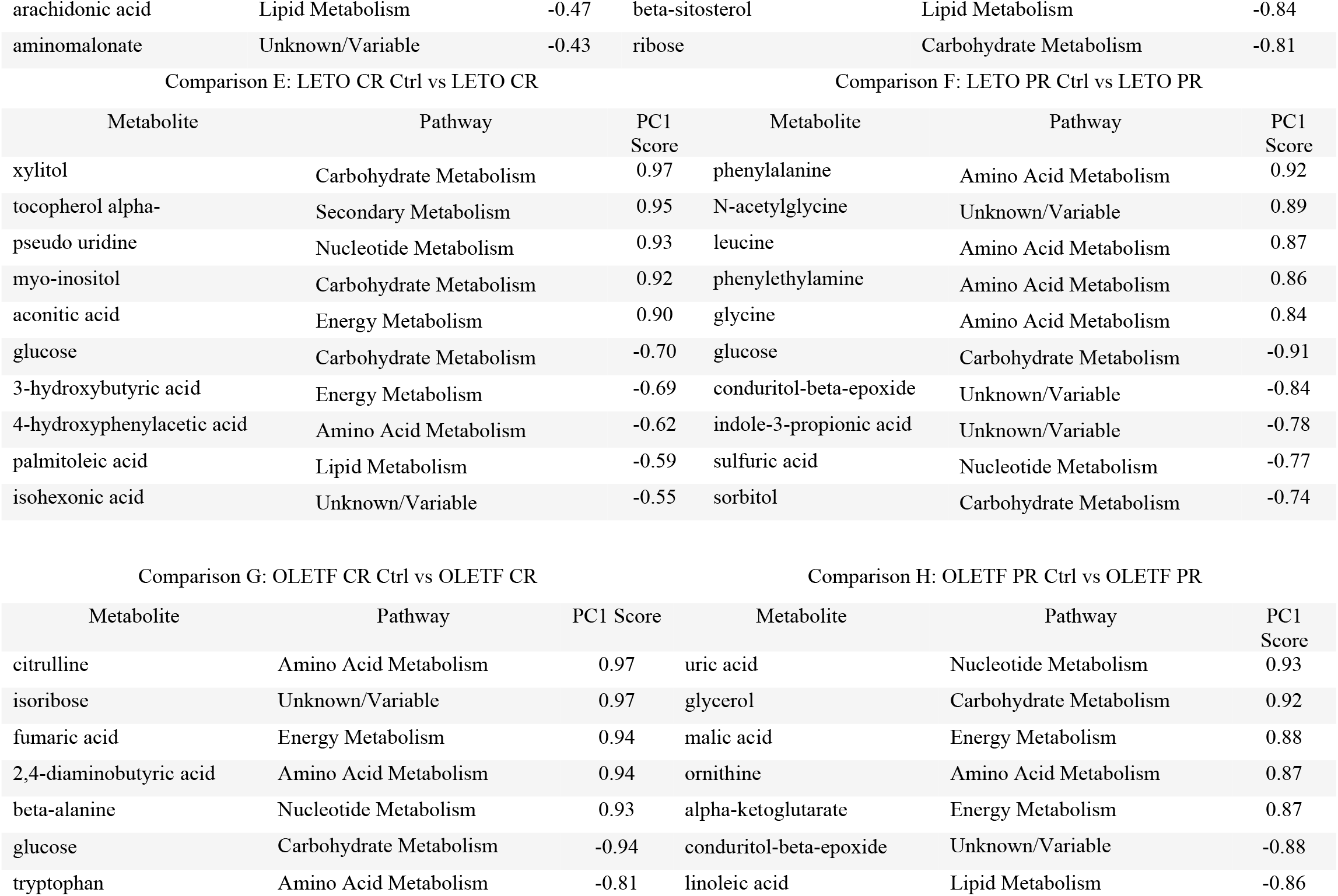
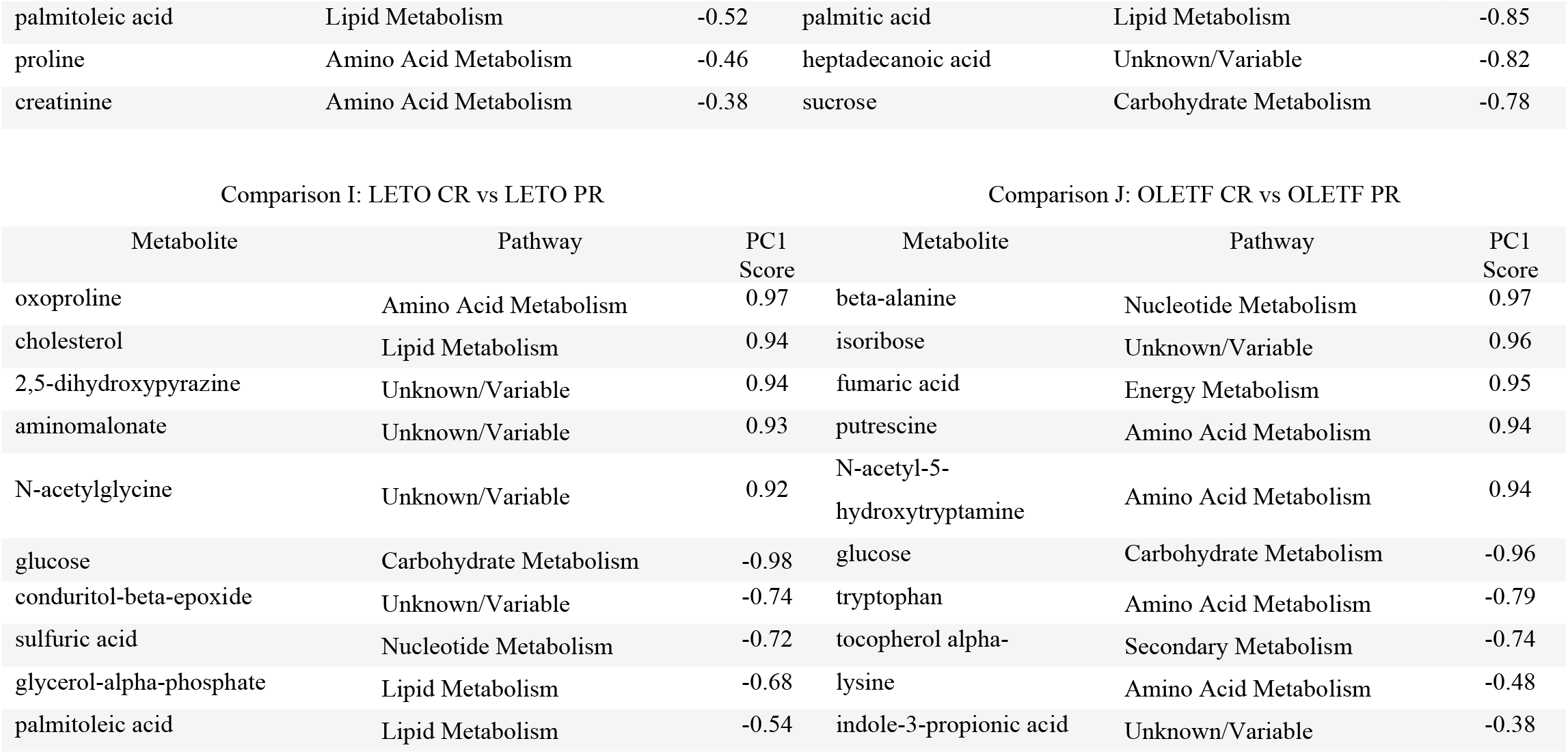
Metabolites with the top 5 positive and negative loadings (by absolute value) for PC1 and their respective pathway classification.

| Comparison A: LETO CR Ctrl vs OLETF CR Ctrl |  |  | Comparison B: LETO PR Ctrl vs OLETF PR Ctrl |  |  |
| --- | --- | --- | --- | --- | --- |
| Metabolite | Pathway | PC1 Score | Metabolite | Pathway | PC1 Score |
| 3-phosphoglycerate | Carbohydrate Metabolism | 0.94 | arachidonic acid | Lipid Metabolism | 0.99 |
| nicotinamide | Secondary Metabolism | 0.93 | threonic acid | Secondary Metabolism | 0.94 |
| threonic acid | Secondary Metabolism | 0.92 | glycine | Amino Acid Metabolism | 0.93 |
| isohexonic acid | Unknown/Variable | 0.91 | serine | Amino Acid Metabolism | 0.93 |
| pseudo uridine | Nucleotide Metabolism | 0.91 | N-acetyl glycine | Unknown/Variable | 0.92 |
| proline | Amino Acid Metabolism | -0.86 | taurine | Lipid Metabolism | -0.89 |
| glucose | Carbohydrate Metabolism | -0.85 | glucose | Carbohydrate Metabolism | -0.87 |
| tocopherol alpha- | Secondary Metabolism | -0.83 | ribitol | Carbohydrate Metabolism | -0.83 |
| ribitol | Carbohydrate Metabolism | -0.78 | hexitol | Carbohydrate Metabolism | -0.83 |
| beta-sitosterol | Lipid Metabolism | -0.78 | ribose | Carbohydrate Metabolism | -0.83 |
| Comparison C: LETO CR vs OLETF CR |  |  | Comparison D: LETO PR vs OLETF PR |  |  |
| Metabolite | Pathway | PC1 Score | Metabolite | Pathway | PC1 Score |
| putrescine | Amino Acid Metabolism | 0.99 | glycerol-alpha-phosphate | Lipid Metabolism | 0.95 |
| 2-deoxypentitol | Unknown/Variable | 0.97 | threonic acid | Secondary Metabolism | 0.94 |
| enolpyruvate | Unknown/Variable | 0.97 | isothreonic acid | Secondary Metabolism | 0.94 |
| pantothenic acid | Amino Acid Metabolism | 0.96 | 3-phosphoglycerate | Carbohydrate Metabolism | 0.92 |
| beta-alanine | Nucleotide Metabolism | 0.95 | pyrophosphate | Energy Metabolism | 0.92 |
| tryptophan | Amino Acid Metabolism | -0.75 | cholesterol | Lipid Metabolism | -0.91 |
| lysine | Amino Acid Metabolism | -0.67 | 2-hydroxybutanoic acid | Secondary Metabolism | -0.88 |
| oxoproline | Amino Acid Metabolism | -0.66 | ribitol | Carbohydrate Metabolism | -0.88 |
| arachidonic acid | Lipid Metabolism | -0.47 | beta-sitosterol | Lipid Metabolism | -0.84 |
| aminomalonate | Unknown/Variable | -0.43 | ribose | Carbohydrate Metabolism | -0.81 |

Comparison E: LETO CR Ctrl vs LETO CR Comparison F: LETO PR Ctrl vs LETO PR
| Metabolite | Pathway | PC1 Score | Metabolite | Pathway | PC1 Score |
| --- | --- | --- | --- | --- | --- |
| xylitol | Carbohydrate Metabolism | 0.97 | phenylalanine | Amino Acid Metabolism | 0.92 |
| tocopherol alpha- | Secondary Metabolism | 0.95 | N-acetylglycine | Unknown/Variable | 0.89 |
| pseudo uridine | Nucleotide Metabolism | 0.93 | leucine | Amino Acid Metabolism | 0.87 |
| myo-inositol | Carbohydrate Metabolism | 0.92 | phenylethylamine | Amino Acid Metabolism | 0.86 |
| aconitic acid | Energy Metabolism | 0.90 | glycine | Amino Acid Metabolism | 0.84 |
| glucose | Carbohydrate Metabolism | -0.70 | glucose | Carbohydrate Metabolism | -0.91 |
| 3-hydroxybutyric acid | Energy Metabolism | -0.69 | conduritol-beta-epoxide | Unknown/Variable | -0.84 |
| 4-hydroxyphenylacetic acid | Amino Acid Metabolism | -0.62 | indole-3-propionic acid | Unknown/Variable | -0.78 |
| palmitoleic acid | Lipid Metabolism | -0.59 | sulfuric acid | Nucleotide Metabolism | -0.77 |
| isohexonic acid | Unknown/Variable | -0.55 | sorbitol | Carbohydrate Metabolism | -0.74 |

Comparison G: OLETF CR Ctrl vs OLETF CR Comparison H: OLETF PR Ctrl vs OLETF PR
| Metabolite | Pathway | PC1 Score | Metabolite | Pathway | PC1 Score |
| --- | --- | --- | --- | --- | --- |
| citrulline | Amino Acid Metabolism | 0.97 | uric acid | Nucleotide Metabolism | 0.93 |
| isoribose | Unknown/Variable | 0.97 | glycerol | Carbohydrate Metabolism | 0.92 |
| fumaric acid | Energy Metabolism | 0.94 | malic acid | Energy Metabolism | 0.88 |
| 2,4-diaminobutyric acid | Amino Acid Metabolism | 0.94 | ornithine | Amino Acid Metabolism | 0.87 |
| beta-alanine | Nucleotide Metabolism | 0.93 | alpha-ketoglutarate | Energy Metabolism | 0.87 |
| glucose | Carbohydrate Metabolism | -0.94 | conduritol-beta-epoxide | Unknown/Variable | -0.88 |
| tryptophan | Amino Acid Metabolism | -0.81 | linoleic acid | Lipid Metabolism | -0.86 |
| palmitoleic acid | Lipid Metabolism | -0.52 | palmitic acid | Lipid Metabolism | -0.85 |
| proline | Amino Acid Metabolism | -0.46 | heptadecanoic acid | Unknown/Variable | -0.82 |
| creatinine | Amino Acid Metabolism | -0.38 | sucrose | Carbohydrate Metabolism | -0.78 |

Comparison I: LETO CR vs LETO PR Comparison J: OLET CR vs OLET PR
| Metabolite | Pathway | PC1 Score | Metabolite | Pathway | PC1 Score |
| --- | --- | --- | --- | --- | --- |
| oxoproline | Amino Acid Metabolism | 0.97 | beta-alanine | Nucleotide Metabolism | 0.97 |
| cholesterol | Lipid Metabolism | 0.94 | isoribose | Unknown/Variable | 0.96 |
| 2,5-dihydroxypyrazine | Unknown/Variable | 0.94 | fumaric acid | Energy Metabolism | 0.95 |
| aminomalonate | Unknown/Variable | 0.93 | putrescine | Amino Acid Metabolism | 0.94 |
| N-acetylglycine | Unknown/Variable | 0.92 | N-acetyl-5-hydroxytryptamine | Amino Acid Metabolism | 0.94 |
| glucose | Carbohydrate Metabolism | -0.98 | glucose | Carbohydrate Metabolism | -0.96 |
| conduitol-beta-epoxide | Unknown/Variable | -0.74 | tryptophan | Amino Acid Metabolism | -0.79 |
| sulfuric acid | Nucleotide Metabolism | -0.72 | tocopherol alpha- | Secondary Metabolism | -0.74 |
| glycerol-alpha-phosphate | Lipid Metabolism | -0.68 | lysine | Amino Acid Metabolism | -0.48 |
| palmitoleic acid | Lipid Metabolism | -0.54 | indole-3-propionic acid | Unknown/Variable | -0.38 |

#### Availability of Data and Material

The datasets generated for this study are available on request to the corresponding author. Raw and normalized metabolomics files can be found in the following repository: https://doi.org/10.6084/m9.figshare.12831005

## Results

### *Ad libitum* fed OLETF rats had lower glycolysis and TCA variables, and higher PPP compared to LETO

OLETF rats had lower basal levels of the glycolysis intermediates, 3-phosphoglycerate (10.9-fold, q<0.001), glyceric acid (2.4-fold, q<0.001), and glycerol-alpha-phosphate (2.5-fold, q=0.001). Moreover, the basal concentrations of citric acid and isocitric acid, the first intermediates of the TCA, were reduced in OLETF compared to LETO (1.9-fold, q=0.032 and 2.0-fold, q=0.061 respectively). Conversely, basal glucose was higher in OLETF (1.3-fold, q=0.020), and ribitol and ribose concentrations were increased in OLETF compared to LETO (2.2-fold, q=0.009 and 1.4-fold, q<0.001, respectively) (**Figure 2A**). These data suggest that during normo-caloric conditions, PPP may contribute significantly to energy production during an insulin resistant condition.

### CR induced higher lipolysis intermediates in OLETF compared to LETO

Compared to LETO after CR, OLETF had higher plasma concentrations of the long-chain fatty acids (LCFA) lignoceric acid (3.1-fold, q=0.087), myristic acid (1.8-fold, q=0.125), and palmitoleic acid (2.6-fold, q=0.038), and the medium-chain saturated fatty acid (MCFA) caprylic acid (2.9-fold, q=0.103), which could reflect a higher rate of lipolysis in OLETF. Moreover, there is an even more pronounced difference between strains in uracil and uric acid concentration (5.3-fold, q=0.041 and 4.9-fold, q=0.021 higher, respectively) (**Figure 2B**), which suggest either an increase in lean tissue catabolism and/or a deficit in kidney function in the insulin resistant strain.

### CR increased intermediates in PPP and TCA, and amino acid concentration, while reducing fasting **glucose in** OLETF

CR in OLETF increased the concentration of the pentoses ribose (2.5-fold, q=0.002) and xylulose (11.6-fold, q=0.003), which are intermediates in the PPP. Moreover, 3-phosphoglycerate, a product of both PPP and glycolysis, increased by 5.6-fold (q=0.009), as well as several intermediates of the TCA cycle: citric acid (2.2-fold, q=0.004), aconitic acid (1.9-fold, q=0.005), isocitric acid (2.2-fold, q=0.008), fumaric acid (5.6-fold, q=0.001), and malic acid (7.2-fold, q=0.001). Furthermore, an increase in the concentration of the amino acids, alanine (1.7-fold, q=0.011), beta-alanine (2.4-fold, q=0.041), cysteine (2.2-fold, q=0.010), serine (1.3-fold, q=0.011), and threonine (1.3-fold, q=0.078), was observed. This translated into a decrease in circulating glucose by 2.6-fold (q<0.001), while increasing adenosine 5-monophosphate (AMP) by 5.2-fold (q=0.005). However, these changes were not observed in LETO after CR. In addition, 5-methoxytrypamine, a metabolite closely related to serotonin, increased 22.3-fold (q=0.038) (**Figure 2C**). The significant increase in plasma AMP and TCA cycle intermediates may reflect an increase in ATP utilization during CR.

### Partial mass recovery increased fasting glucose and decreased TCA, pyrimidine catabolism and lipolysis in OLETF

Partial recovery did not change fasting glucose in LETO, but increased glucose in OLETF by 2.5-fold (q=0.001). Conversely, pyrophosphate, a product of ATP hydrolysis, decreased by 2.3-fold (q=0.007) with an even more pronounced changes in the TCA cycle intermediates, fumaric acid (3.5-fold, q=0.015), malic acid (4.0-fold, q=0.019), and succinic acid (19-fold, q=0.010). Moreover, lignoceric acid (4.4-fold, q=0.010), palmitic acid (2.1-fold, q=0.028), stearic acid (2.5-fold, q=0.017), and caprylic acid (3.7-fold, q=0.006) decreased, along with the pyrimidines, thymine (1.8-fold, q=0.098) and uracil (6.2-fold, q=0.007), its catabolic products, beta-alanine (3.7-fold, q=0.003) and uric acid (4.4-fold, q=0.025), and the glucogenic amino acids, alanine (1.6-fold, q=0.045), cysteine (2.2-fold, q=0.013), glycine (2.5-fold, q<0.001), and aspartic acid (2.0-fold, q=0.016). Interestingly, the concentration of 5-methoxytrypamine was reduced (24.5-fold, q=0.032) after mass recovery in a similar degree as it was increased after CR (**Figure 2D**). Collectively, these changes suggest that, after one week of *ad libitum* diet: **1)** the ATP: ADP ratio is recovered, **2)** hyperglycemia is maintained in OLETF even with decreased endogenous glucose production, and **3)** the reliance on lipid and protein catabolism for energy during insulin resistance is decreased (**Table 4**).

**Table 4.** Main pathway changes after caloric restriction (CR) and partial recovery (PR) compared to ad libitum controls.

|  |  | Gluconeogenesis | Glycolysis | TCA Cycle | PPP | Lipogenesis | Lipolysis | Proteolysis | AA catabolism |
| --- | --- | --- | --- | --- | --- | --- | --- | --- | --- |
| LETO | After CR* | - | - | - | - | - | - | - | - |
|  | After PR** | - | - | - | - | - | - | - | - |
| OLETF | After CR* | ↑ | - | ↑ | ↑ | - | ↑ | ↑ | ↑ |
|  | After PR** | ↑ † | - | - | - | - | ↓ | - | ↓ |

*AA: Amino acid; TCA: Tricarboxylic Acid; PPP: Pentose Phosphate Pathway; \*CR vs CR Ctrl; \*\*PR vs PR Ctrl; ↑ Increase; ↓ Decrease; - no detectable* *change; † shown by previous [18] univariate analysis but not by current metabolomics data*

### Lipid metabolism was inversely correlated to IRI after PR in LETO, but not OLETF

Stearic acid (r=-0.95, q=0.097), palmitoleic acid (r=-0.90, q=0.159), palmitic acid (r=-0.95, q=0.097), myristic acid (r=-0.87, q=0.159), linoleic acid (r=-0.82, q=0.159), glycerol alpha-phosphate (r=-0.94, q=0.097), docohexaenoic acid (r=-0.96, q=0.097), dihydrocholesterol (r=-0.88, q=0.159), cholesterol (r=-0.85, q=0.159), capric acid (r=-0.83, q=0.159), beta-sitosterol (r=-0.83, q=0.159), and arachidonic acid (r=-0.86, q=0.159) had strong, inverse correlations with IRI after PR in LETO. However, no metabolite had a strong, inverse correlation (r<-0.8 and P<0.05) with IRI after CR in LETO nor OLETF, neither after PR in OLETF. (**Figure 3B**). These results suggest that reductions in plasma lipids induced by CR may help maintain normal insulin signaling in LETO, while this protective effect was not observed in the insulin resistant state, and may contribute to the condition.

**Figure 3.**
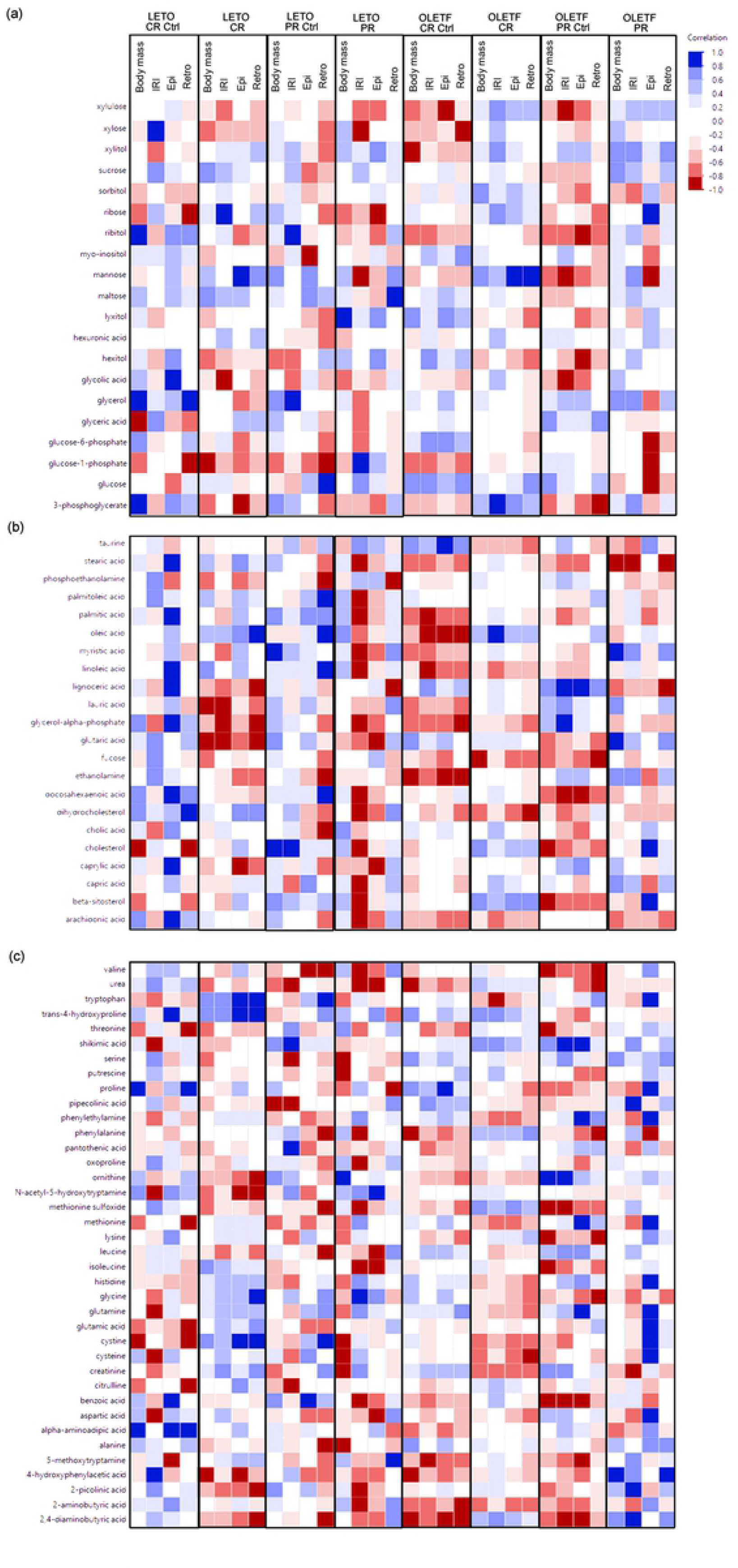
Pearson correlations (r) between Johnson transformed body mass (g), insulin resistance index (IRI) (relative units), retroperitoneal (retro) and epididymal (epi) adipose mass (g) vs Johnson transformed metabolites involved in (a) carbohydrate, (b) lipid and (c) amino acid metabolism.

### Amino acid metabolism was more closely correlated with visceral adiposity than total body mass in OLETF after PR

We previously showed that visceral adipose (retroperitoneal and epididymal) depots were reduced with CR, but depots did not recovered after PR in OLETF suggesting that lean tissue accounted for most of the mass recovery. Moreover, total amino acids increased exclusively in OLETF after CR. Thus, we expected that plasma amino acids would be directly correlated to body mass, rather than adipose, after mass recovery in OLETF. However, contrary to our hypothesis, no amino acid had a strong, positive correlation with body mass, while 4-hydroxyphenylacetic acid (r=0.99, q=0.068) was directly correlated with retroperitoneal fat mass, and tryptophan (r=0.95, q=0.128), phenylethylamine (r=0.91, q=0.172), methionine (r=0.90, q=0.172), histidine (r=0.91, q=0.172), glutamine (r=0.96, q=0.128), glutamic acid (r=0.99, q=0.051), and cystine (r=0.95, q=0.128) were directly correlated with epididymal fat mass in OLETF after PR (**Figure 3C**).

### PLS-DA with subsequent PCA successfully separated each pair of groups at PC1 except for LETO CR vs OLETF CR

PLS-DA was performed for each significant pairwise comparison before performing pairwise PCA analyses (**Figure 4**). Each pairwise comparison had significantly different *(P* < 0.05) PC1 score means, while none of the PC2 score means were significantly different in any of the comparisons. The only exception was comparison C (LETO CR vs OLETF CR) where PC1 scores were not significantly different (**Table 2**). Moreover, this comparison had the closest PC1 score means compared to the rest. This could be explained by the relatively higher variability between samples in the OLETF CR group (SD of 5.80 for PC1). In contrast, LETO CR had relatively much smaller variability, (SD of 0.76 for PC1). Based on these observations, and the fact that PC1 accounted for at least 40% of the variation in all comparisons, further PCA analyses focused on PC1.

**Figure 4.**
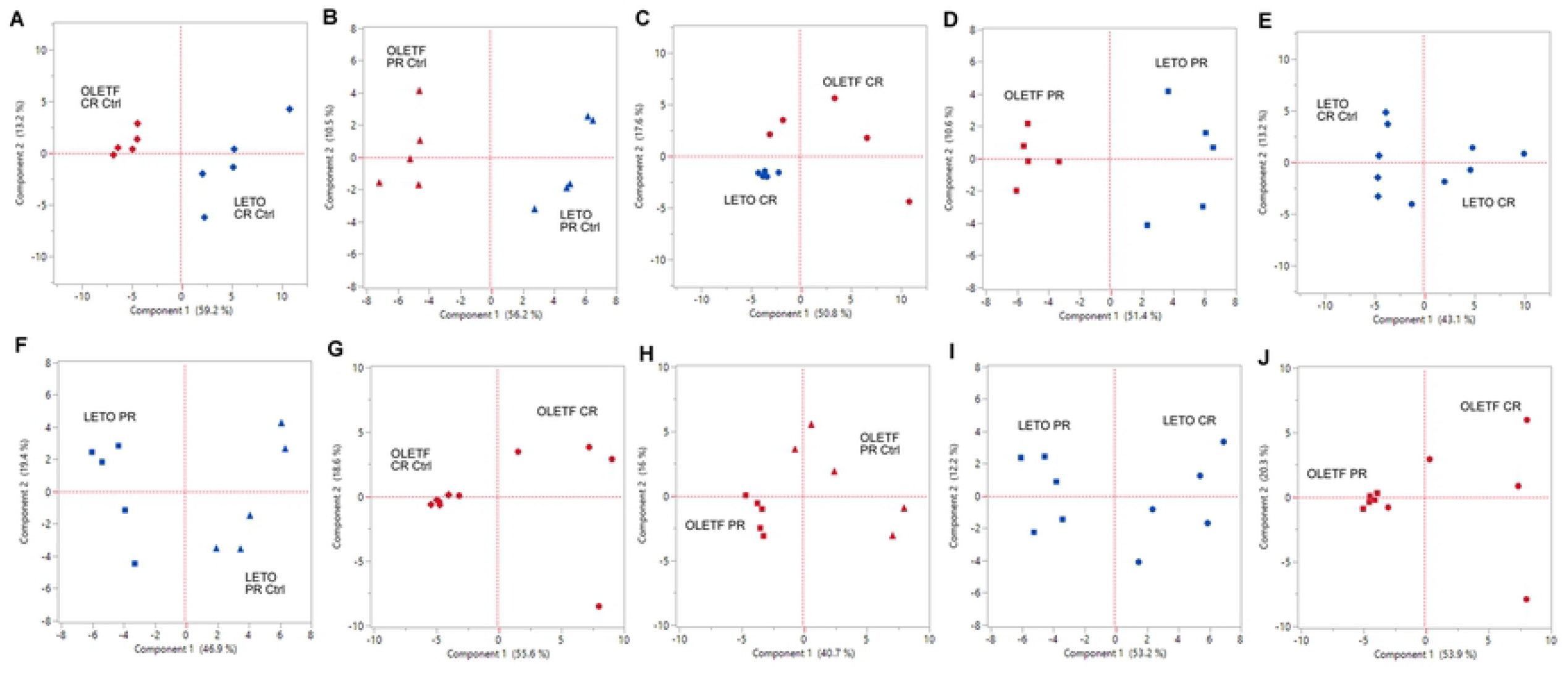
Principal component analysis (PCA) score plots of principal components 1 (x-axis) and 2 (y-axis) for (A) LETO CR Ctrl vs OLETF CR Ctrl, (B) LETO PR Ctrl vs OLETF PR Ctrl, (C) LETO CR vs OLETF CR, (D) LETO PR vs OLETF PR, (E) LETO CR Ctrl vs LETO CR, (F) LETO PR Ctrl vs LETO PR, (G) OLETF CR Ctrl vs OLETF CR, (H) OLETF PR Ctrl vs OLETF PR (I) LETO CR vs LETO PR, and (J) OLETF CR vs OLETF PR.

### Carbohydrate metabolism contributed to the distinction between LETO and OLETF *ad libitum* controls

The monosaccharides, glucose and ribitol, were among the top 5 metabolites that contributed to differentiation between OLETF and LETO (strain effect) on *ad libitum* diet on PC1 at 16 weeks of age (**Table 3, Comparison A**), while hexitol and ribose were added to the list at 17 weeks of age (**Table 3, Comparison B**). Conversely, the metabolites that contributed to the differentiation between the LETO and OLETF CR control groups were involved in lipid, amino acid, and secondary metabolism, with the exception of 3-phosphoglycerate, which is involved in glycolysis. (**Table 3, Comparison A**). These changes could imply a higher reliance in glucose metabolism from the OLETF rats or an impairment in lipid or amino acid metabolism during the insulin resistant state.

### Amino acid metabolism contributed to the distinction between LETO and OLETF after CR, but not after PR

From the top five absolute loadings, 2 metabolites involved in amino acid metabolism (putrescine and pantothenic acid), and beta-alanine, involved in nucleotide metabolism, contributed to the distinction of OLETF from LETO after CR, while the amino acids, tryptophan, lysine, and oxoproline, as well as arachidonic acid (involved in lipid metabolism), contributed to distinguish LETO from OLETF after CR (**Table 3, Comparison C**). This suggests that OLETF rats may rely on protein catabolism in order to synthetize coenzyme A, in contrast to LETO rats that use lipids primarily as a source of energy when carbohydrates are depleted. Conversely, after PR, the metabolites that contributed the most to the distinction between the strains were associated with either lipid metabolism (glycerol-alpha-phosphate for LETO and cholesterol for OLETF) or carbohydrate metabolism (3-phosphoglycerate for LETO, and ribitol and ribose for OLETF) (**Table 3, Comparison D**).

## Discussion

The previous results show that CR elicited more profound metabolic changes in the insulin resistant state compared to the healthy state. Several of the changes observed in glucose metabolism were expected based on previous findings. However, it was rather surprising that these metabolic shifts were not sufficient to spare the utilization of proteins for fuel in the insulin resistant rats, and even more, that mass recovery did not reverted gluconeogenesis to its basal levels, as we will further discuss.

### Carbohydrate metabolism prevailed with an improvement in the TCA cycle, without affecting the oxidative phase of the PPP, after a week of CR in OLETF

A reduction in caloric intake is usually associated with a shift toward increased lipid metabolism because of the reduction in glucose [26-28]. Thus, after 7 days of 50% CR in OLETF rats, we expected to observe a shift toward a greater reliance on lipid metabolism [29]. However, the increases in 3-phosphoglycerate, a by-product of glycolysis, several TCA intermediates, and two intermediates from the non-oxidative phase of the PPP, independent of any detectable changes in oxidative phase intermediates suggest that glucose metabolism is increased with CR during insulin resistance. This highlights the clear benefit of CR to improving glucose metabolism and ameliorating the hyperglycemia during insulin resistance. The increase in plasma ribose likely indicates an increase in PPP concomitant with a decrease on nucleotide synthesis in peripheral tissue [30]. The increase in TCA intermediates suggests that CR even during an insulin resistant state may improve the strain-associated impairment in the TCA cycle. This impairment has been previously described in obese, diabetic rats secondary to hyperphagia and an impairment in the leptin receptor [3], and in T2D patients compared to healthy subjects [31]. The improvement in IRI in OLETF with CR [18] substantiates this data suggesting that CR improves glucose metabolism via enhanced TCA cycle activity. However, in the OLETF the impairment in basal glucose metabolism is likely induced by impaired glycolysis because basal levels of 3-phosphoglycerate were lower in OLETF compared to LETO.

### Partial mass recovery reverted lipolysis, rather than gluconeogenesis, to basal levels

The decrease in glycerol after PR compared to CR suggests that either lipolysis and/or gluconeogenesis is decreased after mass recovery [32], and demonstrates that seven days of partial recovery were sufficient to reverse the limited reliance in lipids in OLETF during an insulin resistant state. This result is consistent with our previous finding where triglyceride levels returned to basal levels following PR [18]. Furthermore, the decrease in the glucogenic amino acids, alanine, cysteine, glycine, and aspartic acid after refeeding (PR), is consistent with the metabolomics data from humans following starvation and refeeding [33]. However, we previously showed strain-dependent changes in the expressions of glucogenic enzymes, although we did not detect consistent changes in these enzymes with CR and PR in OLETF [18] suggesting that the rate of lipolysis, but not gluconeogenesis, is reverted to basal levels in OLETF after PR, Furthermore, comparing our previous measures of hepatic glucogenic enzyme expressions with the current metabolomics data suggests that the lack of detectable changes in protein expression do not accurately reflect the dynamic changes in gluconeogenesis, which puts more of an onus on metabolomics analyses such as these to better complement the molecular data. Additionally, these shifts in lipolysis and gluconeogenesis following PR may highlight the potential detriment of mass recovery, even if only partial as demonstrated here, after CR in an insulin resistant state, as the amelioration of the hyperglycemia with CR may be re-established from the combination of *de novo* glucose synthesis in the liver [34] and impaired glucose uptake from peripheral tissues [35, 36]. If so, this emphasizes the importance of maintaining the reduced body mass after CR as even partial recovery of body mass independent of adiposity has the potential to induce detrimental shifts in glucose metabolism that revert to an insulin resistant, hyperglycemic condition.

### Plasma lipids are better correlated to IRI in a non-insulin resistant state than glucose metabolism intermediates

The lack of an inverse correlation between IR and circulating fatty acids (i.e. products of lipolysis) in an insulin resistant state in the present study may be explained by previous findings where insulin was unable to suppress lipolysis in an insulin resistant state compared to an insulin sensitive condition [37]. This is supported by the lack of changes in plasma lipids in OLETF after CR, even though we previously showed that CR improved IRI in OLETF suggesting that this improvement is largely driven by improvements in glucose metabolism independent of improved lipid metabolism. While an association between long-chain fatty acids and insulin sensitivity in obese, insulin resistant patients has been shown [38] that does not seem to apply during the early onset of insulin resistance in our model.

### Proteolysis is more profound in OLETF than LETO, and is reversed after mass recovery

Normally, during negative caloric imbalance (i.e. CR) most of the amino acids generated after CR are either used by the liver for gluconeogenesis or by the muscle for energy production [39]. Consistent with these findings, we observed small but significant increases in several glucogenic amino acids, which translated into an overall increase in total amino acid concentration after CR in OLETF that returned to baseline levels after PR, but not in LETO. This increase in total amino acids suggests that lean tissue catabolism was increased and that these amino acids may be shuttled into gluconeogenesis to help support the changes in energetic demands. One advantage of this level of CR, however, is that the increased availability of cysteine may increase the production of glutathione [40], thus reducing the overall oxidative damage caused by the insulin resistant state. The potential benefit of PR during an insulin resistant condition is that carbohydrates and lipids return as the primary sources of energy, sparing lean tissue via a reduction in proteolysis. However, the potential detriment of the shift in metabolism is the partial reversal of gluconeogenesis, which could translate into exacerbation of the hyperglycemia in insulin resistant OLETF demonstrating the risk of mass recovery after an acute caloric restriction independent of recovery of visceral adiposity. Furthermore, this highlights the importance of properly managing a caloric restriction regiment following the onset of insulin resistance to ameliorate inappropriately elevated lean tissue catabolism and the potential consequences of this shift in metabolism including an increase nitrogen load on the kidneys [41] and inappropriate muscle loss. These data suggest that a reduction in adipose *per se* may not be sufficient in the long term to improve IRI in an insulin resistant state when gluconeogenesis is inappropriately elevated.

In summary, severe acute caloric restriction was sufficient to elicit significant changes in several metabolic pathways in OLETF but not LETO, and a partial mass recovery comprised mostly by lean tissue reverted most of these changes to baseline levels with the exception of increased gluconeogenesis, and decreased lipolysis and amino acid catabolism. These shifts in substrate metabolism suggest that failure to adhere to this regiment could be detrimental in the long term due to the ability of metabolic shifts to adapt to changes in caloric intake.

## Acknowledgments

We thank J Cazares, B Escobedo, and J Nguyen for their help with the pre-analytical phase of the study.

